# DEEP LEARNING APPROACH FOR RENAL CELL CARCINOMA DETECTION, SUBTYPING, AND GRADING

**DOI:** 10.1101/2024.02.12.577550

**Authors:** Maroof Abdul Aziz, Fatemeh Javadian, Sherin Susheel Mathew, Avinash Gopal, Johannes Stegmaier, Abin Jose

**Affiliations:** Institute of Imaging and Computer Vision, RWTH Aachen University, Germany; Genesys AI Labs, Bangalore, India

**Keywords:** Renal Cell Carcinoma, Pathology, Tumor grading, Hematoxylin and Eosin (H&E) staining, Whole Slide Images

## Abstract

We propose a comprehensive end-to-end pipeline designed for the detection, subtyping, and grading of tumors. Our proposed methodology initiates the generation of a heat map, indicating the severity of the tumor. Subsequently, the identification of the most critical patches is conducted based on the probability scores. These identified patches are then directed to a grade prediction network. A distinctive aspect of our research lies in being the first to explore an end-to-end pipeline for both heat map generation and grading prediction. Our experiments were conducted leveraging the public, The Cancer Genome Atlas (TCGA) repository, focusing specifically on renal cancer. We introduced additional patch-level labels to improve the model performance. The generation of tumor heat maps targeted three primary cancer subtypes: clear cell, papillary, and chromophobe. To enhance our approach, we implemented center-loss and introduced a method aimed at refining the quality of patches. The experimental outcomes highlight superior performance compared to state-of-the-art method. This research contributes to the advancement of tumor detection and grading, emphasizing the significance of an integrated approach for heat map generation and grading pre-diction.

## 1. INTRODUCTION

Cancer [1], a complex group of diseases characterized by the uncontrolled growth and division of abnormal cells, is the focus of this research. Deep learning [2] algorithms have shown remarkable proficiency in analyzing medical images for cancer detection and classification, especially histopathological sections [3], to identify and classify cancerous lesions. This approach is not limited to kidney cancer but includes lung cancer, colorectal cancer, basal cell carcinoma, malignant mesotheliom, breast cancer, prostate cancer, and more [4]. Despite challenges such as the need for manual annotation of large datasets and unknown histopathological features learned by algorithms, the potential of deep learning in pathology is clear [5].

### 1.1. Related work

In the past, there have been several works showing distinguished approaches to using deep learning in medical pathology [6, 7, 8, 9]. Lu et al. proposed in [6] a weakly supervised deep learning approach on Whole Slide Images (WSIs), which is the baseline model used in our work. Further grading of the top-*K* detected regions was inspired by the proposed method in [10, 11]. Similar work is proposed in [12] where a new method to automatically segment nuclei from H&E stained WSIs using convolutional networks [13] is implemented. In [14], deep learning is leveraged to provide Gleason grades [15]. The semi-supervised framework proposed in [16] includes a backbone feature extractor, two task-specific classifiers, and a weight control mechanism. Simultaneous segmentation and classification of nuclei is an efficient method proposed in [17]. The paper [18] proposes a method to overcome the variations in histopathology images that occur due to the nature of the staining procedure, through color deconvolution [19] and color normalization [20]. The work in [3] proposes a patch-level pixel-wise segmentation that was carried out on WSIs with an autoencoder approach. An application of deep learning algorithms on pathologist-level interpretable WSIs is proposed by Zhang et al. in [21]. Liu et al. have proposed in [22] a model that assists breast cancer metastasis detection in lymph nodes. In the literature, most of the works focus either on detection or grading, which motivated us to develop a pipeline to use the most tumorous patches from the heat map prediction model to a network that predicts the grades.

### 1.2. This paper

This paper mainly deals with the nuances of cancer in the specific context of kidney pathology [23] and focuses on three major sub-types, namely, clear cell renal cell carcinoma (ccRCC) [24], papillary renal cell carcinoma (pRCC) [25], and chromophobe renal cell carcinoma (chRCC) [26]. We use a combination of weakly supervised and supervised training, coupled with center loss [27], to train an attention model and generate tumor heat maps. Once the heat maps are generated, the top-*K* patches are passed to the grading module, which predicts the grade. The proposed approach is explained in Section 2. Training details are elaborated in Section 3. Datasets, experimental findings, and limitations of our method are discussed in Section 4. Concluding remarks and future work are discussed in Section 5.

## 2. PROPOSED APPROACH

In this paper, we introduce a comprehensive end-to-end pipeline designed for the detection, subtyping, and grading of Renal Cell Carcinoma (RCC) using WSIs. Fig. 1 shows the different steps involved in the detection and grading. Initially, the Hematoxylin and Eosin (H&E) stained WSIs undergo a segmentation process to eliminate empty or non-informative regions. Subsequently, these images are subdivided into smaller, manageable patches. Each patch is then processed through a CNN-based encoder to extract its feature embeddings. These embeddings are subsequently passed through an attention network, which assigns an attention score to each patch, reflecting its relevance to the identified class. This process results in patch-level features, each weighted by its corresponding attention score, culminating in a comprehensive slide-level representation for accurate classification. Furthermore, the model generates heat maps derived from these attention scores, offering a visual representation of the tumor regions. Patches receiving the highest attention scores are then selectively advanced to the subsequent phase of our pipeline, dedicated to the precise grading of RCC. The foundational architecture of our model is derived from the CLAM model [6]. We have expanded upon this network, integrating several key improvements detailed in the following sections.

**Fig. 1:**
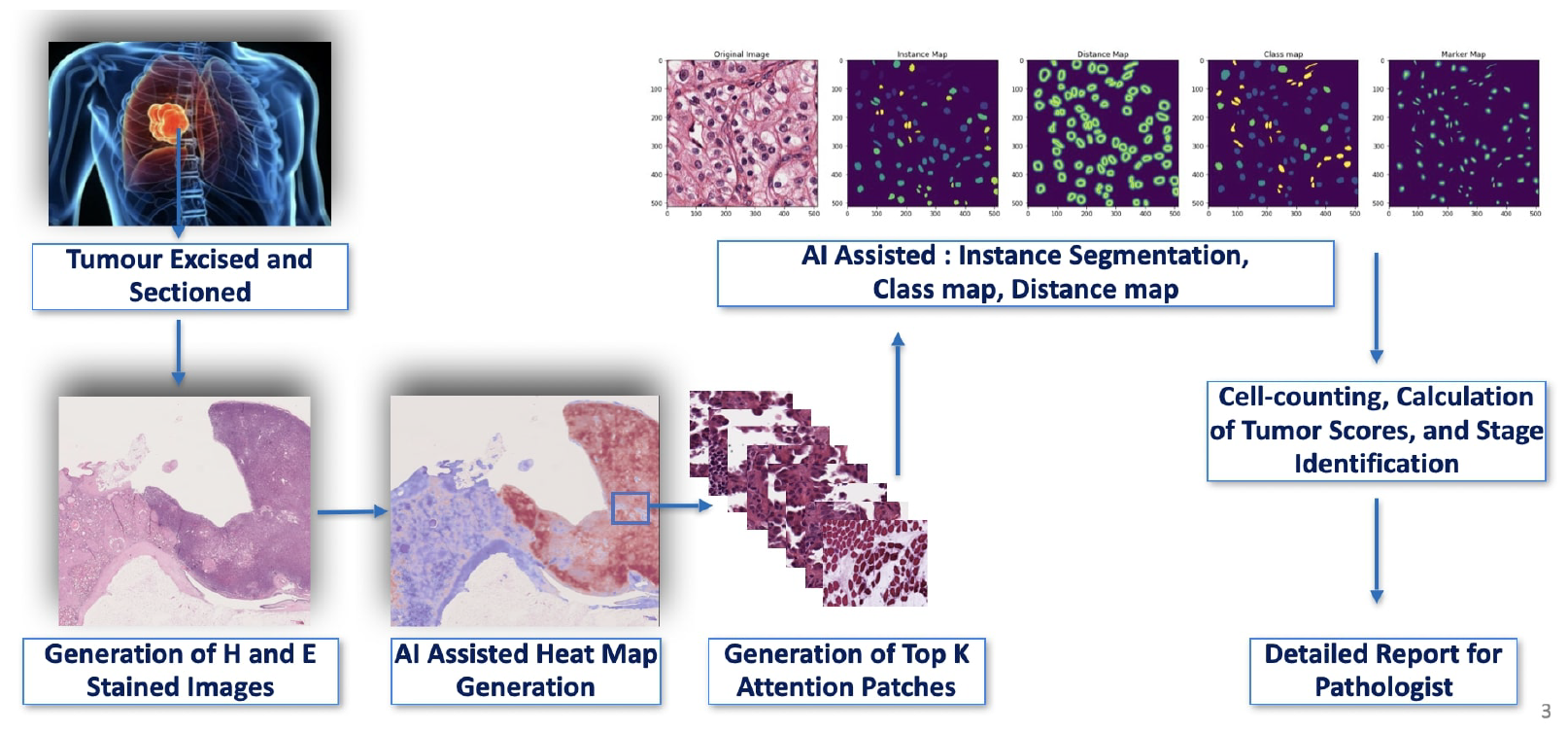
Block diagram of the proposed end-to-end pathology pipeline. 1. In the first step, H and E stained images are taken from the WSI scanner and the heat map image is generated which shows the tumor probability, with red showing higher tumor probability. The next step is the nuclei grading. Here, the instance map, distance map and class maps are generated. The grading of the tumor is done in this step for the top-*K* patches.

**Fig. 2:**
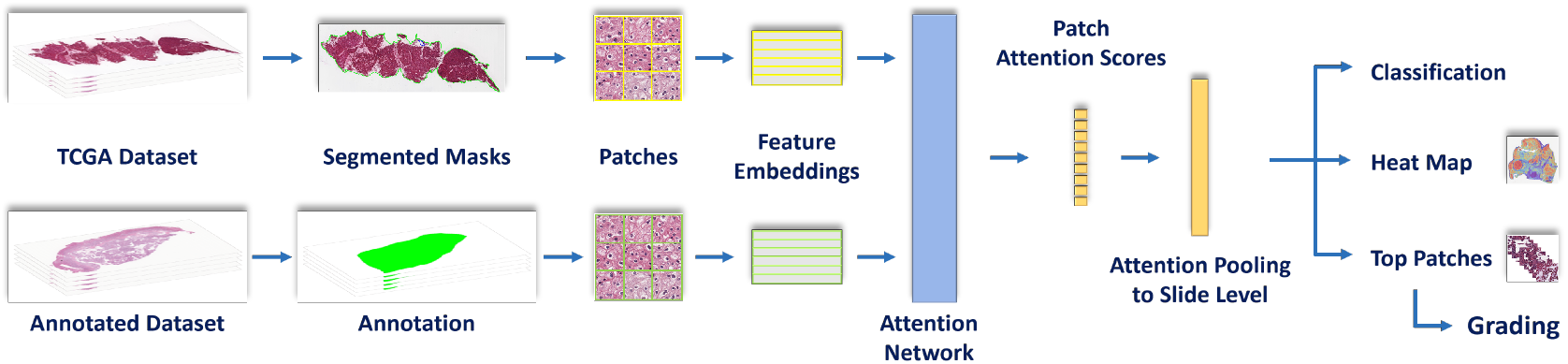
Integration of slide-level and patch-level annotated datasets. TCGA dataset undergoes segmentation to produce segmented masks, while the annotated dataset provides precise patch-level labels. Both datasets contribute patches that are processed to extract feature embeddings. These embeddings are fed into an attention network that yields patch attention scores. Attention pooling aggregates these scores to a slide-level representation for classification. The output includes a heat map for visual analysis and the selection of top-*K* patches, which are then forwarded for detailed grading of RCC.

### 2.1. Segmentation and patch generation

This paper features an improved segmentation method, as illustrated in Fig. 3, designed to minimize the inclusion of white patches in the training process. Initially, the WSIs are segmented using a classical contour-based approach. Subsequently, an additional filtering step is employed to remove patches that predominantly contain white regions, which are typically empty areas. This refinement is aimed at mitigating potential noise and inaccuracies such white patches could introduce, thereby ensuring a more refined training dataset for the model. Additionally, considering TCGA dataset ^1^ contains WSIs in both 40x and 20x magnifications, we have implemented a systematic approach to normalize the viewing area. Such standardization ensures that the model processes patches corresponding to consistent physical tissue sizes, irrespective of the original magnification of the WSI, thereby maintaining uniformity and accuracy in the detection and analysis process. This is achieved by adjusting the size of the patches and employing downsampling as shown in Fig. 4.

**Fig. 3:**
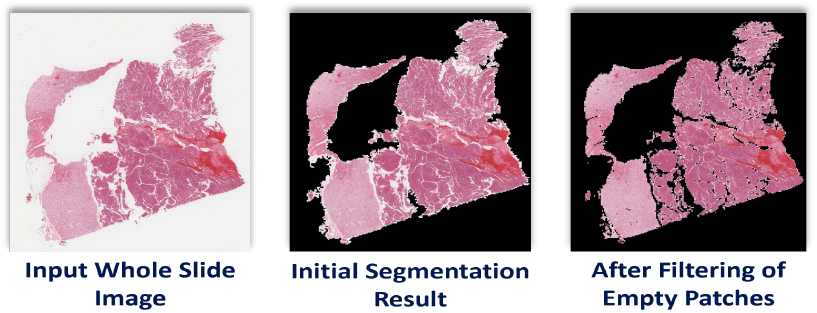
Segmentation improvement through filtering of empty patches. The input WSI is first segmented using a simple contour segmentation, after which patches with a majority of white regions are further filtered out.

**Fig. 4:**
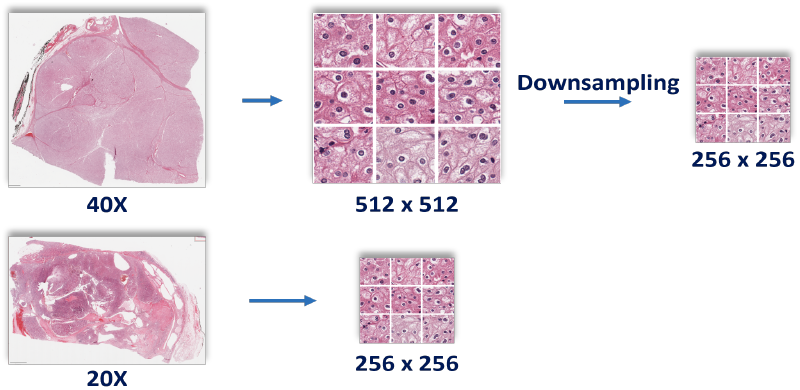
Uniform viewing area irrespective of magnification. WSIs with 20x magnification are directly patched at 256 *×* 256 pixel size while those with 40x magnification are first patched at 512 *×* 512 pixel size and then downsampled to 256 *×* 256 size.

### 2.2. Feature extraction and attention network

Following the patch extraction, feature embeddings for each patch are computed using a ResNet-50 [28] model pre-trained on the ImageNet dataset [29]. These embeddings are then channelled into an attention network as described by Lu et al. [6], which assigns attention scores to each patch, based on its relevance to each class under consideration. The patch feature embeddings are then weighted according to their respective attention scores and aggregated to construct a cohesive slide-level representation, as shown in Fig. 2. Furthermore, the model employs the computed attention scores to generate detailed heat maps, which visually highlight tumorous regions within the slides with gradients of blue indicating non-tumorous and gradients of red indicating tumorous regions. These maps, alongside the identification of top-*K* patches based on attention scores, provide a granular view of the tissue characteristics, aiding in the interpretability of the model’s predictions. The top-*K* patches of tumorous WSIs are then passed on to the grading part of the pipeline.

### 2.3. Incorporation of center-loss

We have incorporated center loss into the total loss function to further refine the model’s performance by effectively reducing the intraclass variance and increasing the inter-class variance at the patch-level. The intra-class center loss (*L*_center,intra_) is the mean squared error between the feature vectors **h** and the corresponding class centers **c**_*i*_, computed over *N* patches as given below:

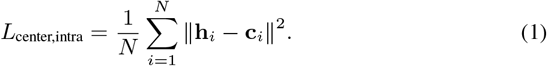

The inter-class center loss (*L*_center,inter_) is computed based on the Euclidean distances between each pair of class centers *d*(**c**_*i*_, **c**_*j*_) for all *M* classes as defined below:

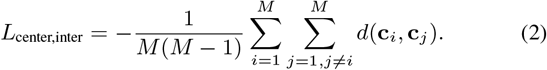

The total center loss (*L*_center,total_) is a weighted sum of the intra-center and inter-center losses, with the hyperparameter, *α* controlling the trade-off between both and is defined as, *L*_center,total_ = *α · L*_center,intra_ + (1 *− α*) *· L*_center,inter_. While the primary classification loss focuses on the correct labelling at the slide-level, center loss provides an additional criterion focused on the geometric distribution of features at the patch-level, leading to more accurate heat maps.

### 2.4. Incorporation of annotated dataset for patch-level labels

This work introduces a hybrid approach as shown in Fig. 2, by combining a slide-level labelled dataset from TCGA for weakly supervised learning, with a dataset featuring annotation masks [10] to extract patch-level labels. This methodology significantly improves the accuracy of heat map generation and top-*K* patch selection for precise grading post-classification.

#### Loss calculation for WSIs with only slide-level labels

Patches extracted from images possessing only slide-level labels, inherit the label of its parent slide (weakly supervised learning). For these images, the loss is computed as a combination of classification loss, instance-level clustering loss, and center loss. Instance-level clustering loss *L*_inst_, as described by Lu et al. [6], leverages pseudo labels from attention network outputs to cluster patches within a WSI, enhancing feature discrimination by separating highly attended patches as class-specific evidence from less attended patches as negative evidence. The initial loss for slide-level labeled images (*L*_slide,init_) is the sum of slide-level cross entropy classification loss (*L*_class_) and instance-level clustering loss (*L*_inst_), with a regularisation factor *λ*_inst_ as given by, *L*_slide,init_ = *λ*_inst_ *· L*_inst_ + (1 *− λ*_inst_) *· L*_class_. The initial slide loss is then combined with the center loss (*L*_center,slide_) with regularisation factor *λ*_center_ to get the total loss for slide-level labeled images (*L*_slide,total_) and is defined as, *L*_slide,total_ = *λ*_center_ *· L*_center,total_ + (1 *− λ*_center_) *· L*_slide,init_.

#### Loss calculation for WSIs with patch-level labels

Conversely, patches derived from images with detailed annotation masks are labelled in accordance with the specific regions highlighted by these masks. For these patches, the loss is calculated as a combination of cross entropy classification loss (*L*_class_) and center loss (*L*_center,patch_) with regularisation factor *λ*_center_, which is expressed as, *L*_patch,total_ = *λ*_center_ *· L*_center,total_ + (1 *− λ*_center_) *· L*_class_.

#### Final loss calculation

The final loss for model training (*L*_final_) is a combination of losses of both slide-level and patch-level annotated datasets computed in the previous steps. It is fine-tuned by allocating additional weightage *λ*_patch_ to the classification of images sourced from the patch-level annotated dataset, to ensure that the model effectively leverages the detailed annotations available, leading to more accurate and granular learning as defined as, *L*_final_ = *λ*_patch_ *· L*_patch,total_ + (1 *− λ*_patch_) *· L*_slide,total_.

### 2.5. Evaluation criteria and hyperparameter tuning

#### Evaluation criteria

The efficacy of our model in the classification of RCC WSIs is assessed using a comprehensive set of metrics which include accuracy, Balanced Accuracy (bACC), Area Under Curve (AUC), F1 score, and Mean Average Precision (mAP). For patch-level analysis, quantifying accuracy presents unique challenges, as this level of granularity is not easily captured through standard metrics without ground truth. Therefore, the accuracy at the patch-level is primarily evaluated based on qualitative feedback from expert pathologists. This approach ensures a practical and real-world assessment of the model’s performance. Additionally, the generation of top-*K* patches for the grading pipeline is an integral part of our evaluation, as these patches are crucial for precise tumor grading.

#### Hyperparameter tuning

The tuning of hyperparameters is pivotal in optimizing our model’s performance. The key hyperparameters tuned in our work include learning rate, weight decay, regularization coefficients for instance-level clustering loss (*λ*_inst_), center loss (*λ*_center_), and the patch-level annotated dataset (*λ*_patch_). The judicious selection and tuning of these hyperparameters are critical in achieving the desired balance between learning efficiency and model robustness, ultimately enhancing the overall performance.

### 2.6. Grading network

In our paper, we utilize the HRFE model [11] as a crucial component in RCC grading, marking a significant step in computational pathology. The process begins with the identification of key tumorous regions, where the top-*K* patches from the tumor heat map, generated in a preceding step, are fed into this grading module [10]. This important step ensures that only the most relevant sections of the tissue receive detailed analysis. The model’s segmentation phase is motivated by creating distance and binary maps from [30], crucial for distinguishing each nucleus from the surrounding tissue. This segmentation is decisive for the subsequent grading, where each identified nucleus is precisely analyzed. To handle inter-class similarity property, the model employs two magnification levels to classify nucleoli grading [31]. To evaluate its effectiveness, an extensive test database of unannotated images was utilized. The model produces overlay images based on its predictions, which are subsequently merged with the original images, enhancing the visual representation for more detailed analysis. These composite images are presented to pathologists for a human validation process, enabling a thorough assessment of the model’s accuracy.

## 3. TRAINING DETAILS

### 3.1. Baseline model

The foundational step in our model’s development involved establishing a baseline for comparison. To this end, we trained the CLAM [6] model to classify WSIs from TCGA renal dataset into four classes: Normal tissues, ccRCC, pRCC, and chRCC. The dataset comprised 1180 WSIs, distributed as follows: 300 Normal, 400 ccRCC, 300 pRCC, and 180 chRCC. These were divided into training, validation, and test sets in an 80:10:10 ratio, respectively. This baseline model served as a benchmark against which the performance of our proposed methodology could be measured.

### 3.2. Proposed model

Building upon the baseline, our proposed model introduces a more intricate approach by integrating a patch-level annotated dataset into the training process along with center loss implementation and improved segmentation. This dataset complements the existing TCGA renal dataset and brings a higher resolution of detail to the training phase, allowing for a more nuanced differentiation between the four cancerous and non-cancerous classes at the patch-level. For the proposed model, the same validation and test datasets used in the baseline model were employed to maintain consistency in performance evaluation. However, the training set was expanded to include the additional patch-level annotated WSIs, distributed as follows: 100 ccRCC, 100 pRCC, and 34 chRCC. The extended training dataset ensures that the model not only learns from the global slide-level labels but also gains insights from localized annotations which would be crucial for heat maps and top-*K* patches for the grading pipeline. The source code is available on GitHub^2^.

### 3.3. Training the grading model

The incorporation of a fine-grained classification network [32] for each RCC nucleus, unlike previous methods that separate segmentation and classification processes [33], results in higher performance with a less complex model. Current advancements [17] employ unified frameworks to enhance both tasks so the classification branch is integrated with a segmentation network. This is achieved by using high-resolution feature extractors and a two-stage learning framework [34]. The current applied network for grading [11] justify the complexities of tumor nuclei grading in histopathological images by combining advanced segmentation and feature extraction. The dataset, comprising ccRCC and pRCC H&E stained images from the TCGA repository, utilizes 90% of its images for model training and validation. The remainder is used for evaluating the model’s numerical accuracy, supplemented by top-*K* unannotated images from the preceding network.

## 4. EXPERIMENTAL RESULTS

### 4.1. Detection and subtyping

The performance metrics presented in Table 1 showcases the model’s improved performance gains over the baseline in tumor subtyping. With the integration of center loss and annotated data, the increase in AUC, bACC, mAP, and F1 score signifies a more precise classification capability. The amelioration in feature space clustering is evident in Fig. 6

**Table 1:** Slide-level performance metrics compared to the baseline [6].

| Model | AUC | bACC | mAP | F1 |
| --- | --- | --- | --- | --- |
| Baseline [6] | 0.9835 | 0.8985 | 0.8372 | 0.8973 |
| Ours | **0.9862** | **0.9062** | **0.8466** | **0.9057** |

**Fig. 5:**
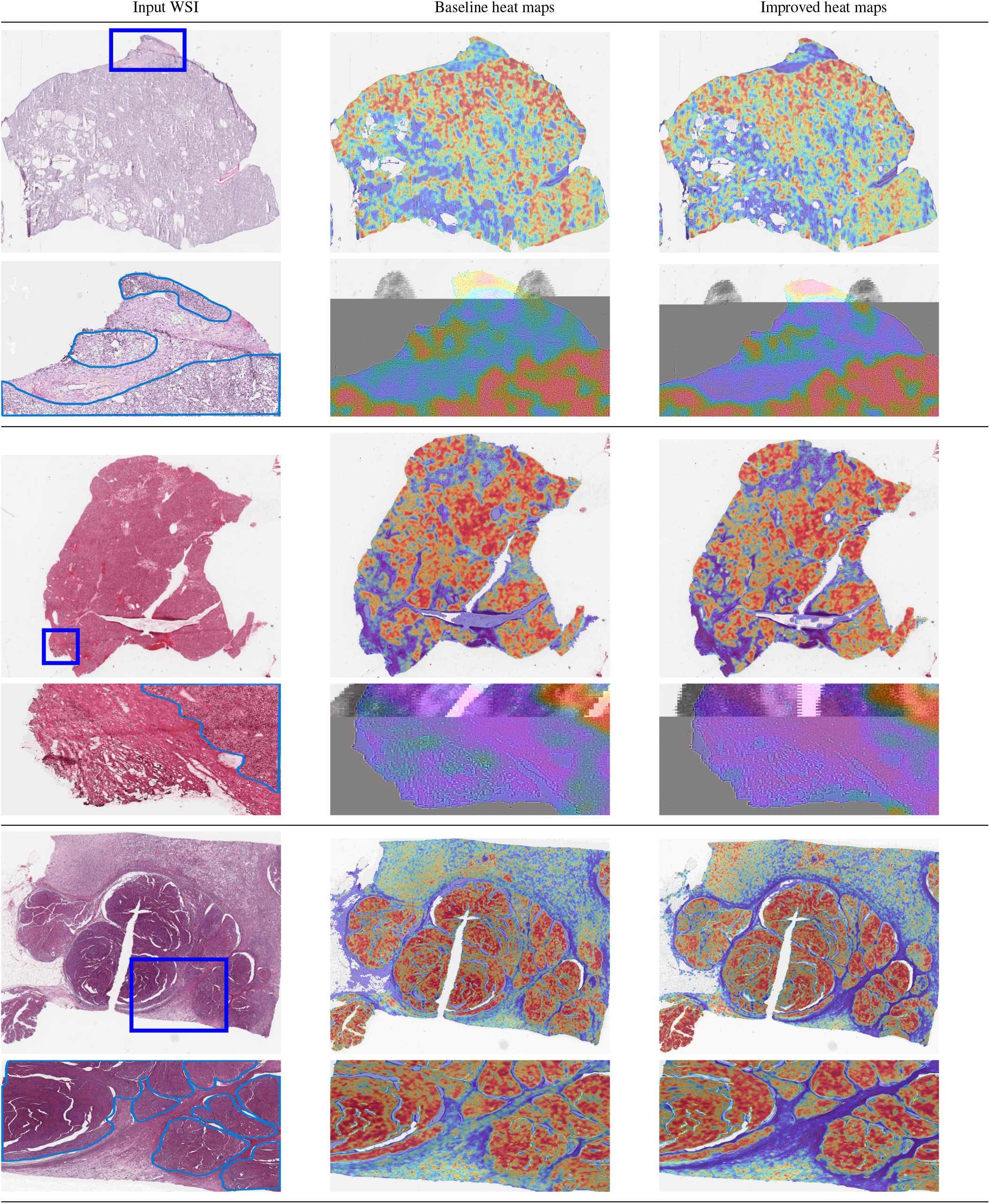
Column 1 presents the original HE stained WSIs alongside pathologists’ annotations. Column 2 shows the heat map results from the baseline model, depicting the initial differentiation between tumor and normal tissue. Column 3 displays the results from the improved model, demonstrating enhanced discrimination of tumor (depicted in red) versus normal tissue (depicted in blue) and the exclusion of non-relevant areas such as fatty tissues and empty spaces. This reflects a more accurate alignment with the pathologists’ annotations. In each result, the first row indicates the WSI, and the next row indicates the zoomed region, with the blue box in the original WSI indicating the region that is zoomed.

**Fig. 6:**
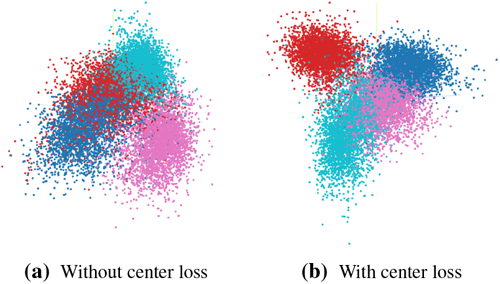
t-SNE [35] plot indicating the compactness in feature space by the addition of center loss. With center loss, the intra-class scatter has reduced as shown in (b). The four classes are normal patches (pink), clear-cell (cyan), papillary (red), and chromophobe (blue), RCCs.

As shown in Fig. 5, the enhanced model exhibits a marked refinement in differentiating tumor regions (red) from normal tissue (blue) when compared to the baseline model. Moreover, the model demonstrates a refined heat map generation of the tissue composition by specifically discounting fatty tissues and empty regions, which are often present in WSIs. These heat maps exhibit an increased level of confidence, providing clear demarcations between the pathological and non-pathological areas within the WSIs. These enhancements reflect a significant step forward in the precise classification of WSIs for renal carcinoma, offering a more dependable diagnostic aid.

### 4.2. Grading

The grading network is fed with 40x magnified patches selected as the top-*K* patches from the tumor heat map, previously identified in an earlier stage of our pipeline. This network has demonstrated its capability to accurately grade both ccRCC and pRCC in H&E stained images. The grading of nuclei in both pRCC and ccRCC depends on certain criteria, including the size and prominence of nucleoli and the nucleocytoplasmic ratio. Within the tumor regions of ccRCC and pRCC, two types of nuclei are present – endothelial and tumor nuclei, ranging from grade 1 to 3. In the classification map, our approach incorporates majority voting to determine the most prevalent class. Despite the model’s competence in generating instance maps, which include accurately segmenting nuclei even when overlapping or adjacent, it encounters challenges in differentiating close grades, such as grades 1 and 2, or 2 and 3. We explored the efficacy of grayscale H channel in transmittance and absorbance modes, compared to conventional RGB H&E images for grading [18]. Our analysis covers a variety of metrics including the Aggregated Jaccard Index (AJI), Panoptic Quality (PQ), Precision, Recall, and F1 scores as depicted in Table 2.

**Table 2:** Nuclei grading across different imaging modes.

| Metric | H&E | H trans. | H absorb. |
| --- | --- | --- | --- |
| AJI | **0.7349** | 0.7265 | 0.7234 |
| Mean PQ | **0.5272** | 0.5016 | 0.4990 |
| Overall F1 | **0.7474** | 0.7275 | 0.7324 |
| Grade 1 F1 | **0.8243** | 0.8113 | 0.8151 |
| Grade 2 F1 | **0.4793** | 0.4290 | 0.4403 |
| Grade 3 F1 | **0.5228** | 0.5189 | 0.5229 |
| Endothelial F1 | **0.7057** | 0.6761 | 0.6729 |

The results depicted in Fig.7 indicated that while the H&E images outperformed in nearly all metrics, there wasn’t a clear winner between the H channel absorbance and transmittance modes. Each mode excelled in certain aspects but fell short in others. For instance, the H channel absorbance mode showed higher performance in some metrics like overall F1 and F1 particularly for certain grades, while the transmittance mode performed better in AJI, PQ and endothelial F1 scores. However, both modes exhibited lower scores in critical areas such as PQ for grades 2, 3, and endothelial compared to H&E images, highlighting their limitations in accurate classification of these specific grades. The original H&E images remained the superior method in our study. Beyond classification and heat map generation, the model generates top-*K* patches based on the highest attention scores. When the classification outcome indicates ccRCC or pRCC, these top-*K* patches serve as a critical input for the subsequent grading process, ensuring that the grading phase is informed by patches that exhibit the strongest indications of malignancy. Fig. 8 indicates the grades predicted by the network.

**Fig. 7:**
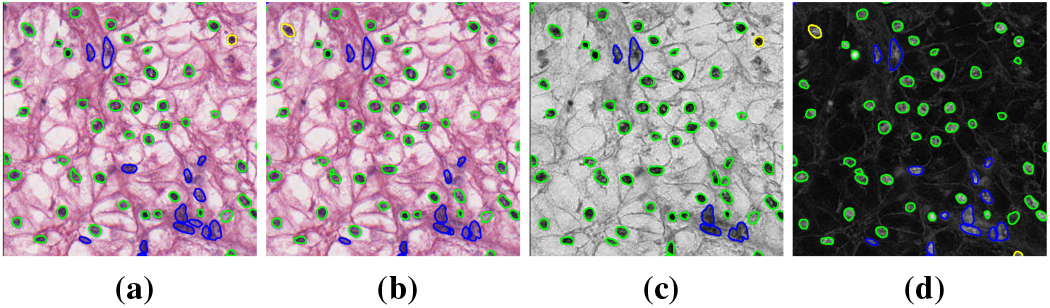
Ground-truth (a), Prediction for H&E (b), Transmittance H Channel (c) and Absorbance H Channel (d).

**Fig. 8:**
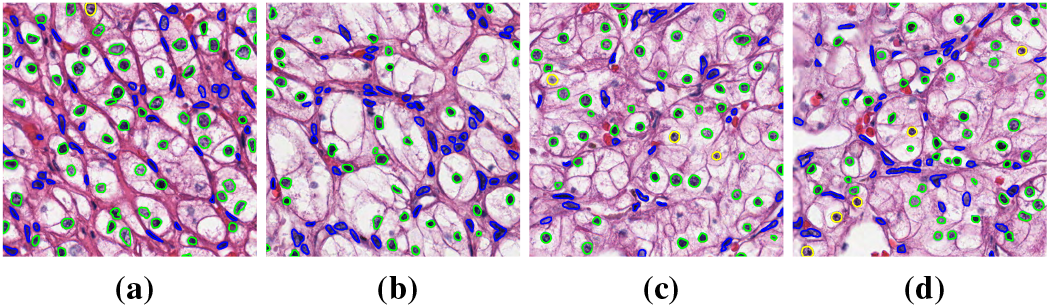
Grade generated by the network (Yellow: grade 2, Green: grade 1, Blue: Endothelial)

### 4.3. Limitations and areas to improve

While our model marks advancement in RCC classification and grading, few areas for enhancement have been identified to refine its diagnostic accuracy. Challenges include the model’s occasional misclassification of blurred tumorous regions and normal tissues (Fig. 9), indicating a need for further improved recognition of subtle pathological features. Additionally, the limited diversity of RCC subtypes in the TCGA dataset restricts the model’s comprehensiveness. Addressing these issues by enhancing the model’s interpretative capabilities and expanding the dataset will be pivotal in elevating the model’s performance and its applicability across a wider spectrum of RCC variants.

**Fig. 9:**
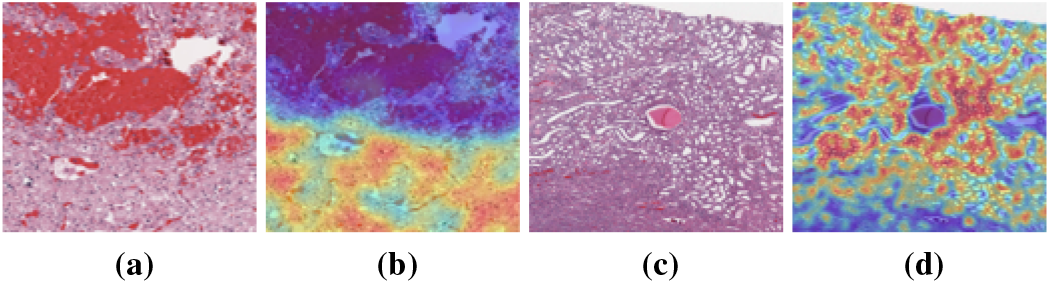
WSI results where our algorithm has limitations. (a) WSI containing tumorous blurred regions. (b) model is not able to classify the blurred regions properly. (The natural blur introduced during WSI acquisition causes this problem). (c) WSI patch containing normal tissues (d) model miss classifies, few normal patches as tumorous patches.

## 5. CONCLUSIONS

In conclusion, our proposed pipeline for tumor detection and grading has demonstrated remarkable effectiveness, surpassing the performance of existing state-of-the-art methods. The inclusion of a heat map generation step, coupled with the identification of tumor grades from the most informative patches, underscores the robustness of our approach. Our experimentation, conducted on the TCGA repository for renal cancer, specifically addressed three distinct cancer subtypes: chromophobe, papillary, and clear cell carcinomas. The integration of center-loss and the introduction of a patch enhancement method significantly contributed to the overall improvement of the quantitative measures compared to the baseline model, and enhanced the demarcation between tumor and non tumor patches. Our future research endeavours will focus on refining the heat map prediction pipeline through knowledge distillation to achieve more accurate patch generation during the training phase. The validation of our findings on real datasets holds promise for providing additional insights and solidifying the practical applicability of our proposed pipeline. Moreover, we aspire to enhance the grading module further, tailoring it specifically for chromophobe and papillary carcinomas. By expanding our experimentation to these specific cancer subtypes, we aim to gain deeper insights into the nuanced characteristics and intricacies associated with each subtype.

## Footnotes

1 https://www.cancer.gov/tcga

2 https://github.com/RoboMaroof/Cancer-Detection-on-WSIs.git

## REFERENCES

[1] Motzer et al., “Renal-cell carcinoma,” New England Journal of Medicine, vol. 335, no. 12, pp. 865–875, 1996.

[2] LeCun et al., “Deep learning,” Nature, vol. 521, no. 7553, pp. 436–444, 2015.

[3] Wang et al., “Pathology image analysis using segmentation deep learning algorithms,” The American Journal of Pathology, vol. 189, no. 9, pp. 1686–1698, 2019.

[4] Ali et al., “State-of-the-art challenges and perspectives in multi-organ cancer diagnosis via deep learning-based methods,” Cancers, vol. 13, no. 21, pp. 5546, 2021.

[5] Hägele et al., “Resolving challenges in deep learning-based analyses of histopathological images using explanation methods,” Scientific Reports, vol. 10, no. 1, pp. 6423, 2020.

[6] Lu et al., “Data-efficient and weakly supervised computational pathology on whole-slide images,” Nature Biomedical Engineering, vol. 5, no. 6, pp. 555–570, 2021.

[7] Farzad et al., “Digital imaging in pathology: whole-slide imaging and beyond,” Annual Review of Pathology: Mechanisms of Disease, vol. 8, pp. 331–359, 2013.

[8] Liron et al., “Review of the current state of whole slide imaging in pathology,” Journal of Pathology Informatics, vol. 2, no. 1, pp. 36, 2011.

[9] Neofytos Dimitriou, Ognjen Arandjelovicć, and Peter D Caie, “Deep learning for whole slide image analysis: an overview,” Frontiers in Medicine, vol. 6, pp. 264, 2019.

[10] Gao et al., “Renal cell carcinoma detection and subtyping with minimal point-based annotation in whole-slide images,” in International Conference on Medical Image Computing and Computer-Assisted Intervention. Springer, 2020, pp. 439–448.

[11] Gao et al., “Nuclei grading of clear cell renal cell carcinoma in histopathological image by composite high-resolution network,” arXiv preprint arXiv:2106.10641, June 2021.

[12] Peter Naylor, Marick Laé, Fabien Reyal, and Thomas Walter, “Segmentation of nuclei in histopathology images by deep regression of the distance map,” IEEE Transactions on Medical Imaging, vol. 38, no. 2, pp. 448–459, 2019.

[13] Sultana et al., “Evolution of image segmentation using deep convolutional neural network: A survey,” Knowledge-Based Systems, vol. 201, pp. 106062, 2020.

[14] Madabhushi et al., “Deep-learning approaches for gleason grading of prostate biopsies,” The Lancet Oncology, vol. 21, no. 2, pp. 187–189, 2020.

[15] Ni Chen and Qiao Zhou, “The evolving gleason grading system,” Chinese Journal of Cancer Research, vol. 28, no. 1, pp. 58, 2016.

[16] Gao et al., “A semi-supervised multi-task learning framework for cancer classification with weak annotation in whole-slide images,” Medical Image Analysis, vol. 83, pp. 102652, 2023.

[17] Graham et al., “Hover-net: Simultaneous segmentation and classification of nuclei in multi-tissue histology images,” Medical Image Analysis, vol. 58, pp. 101563, 2019.

[18] Bianconi et al., “Experimental assessment of color deconvolution and color normalization for automated classification of histology images stained with hematoxylin and eosin,” Cancers, vol. 12, no. 11, pp. 3337, 2020.

[19] Arnout C Ruifrok, Dennis A Johnston, et al., “Quantification of histochemical staining by color deconvolution,” Analytical and Quantitative Cytology and Histology, vol. 23, no. 4, pp. 291–299, 2001.

[20] Devrim Onder, Selen Zengin, and Sulen Sarioglu, “A review on color normalization and color deconvolution methods in histopathology,” Applied Immunohistochemistry & Molecular Morphology, vol. 22, no. 10, pp. 713–719, 2014.

[21] Zhang et al., “Pathologist-level interpretable whole-slide cancer diagnosis with deep learning,” Nature Machine Intelligence, vol. 1, no. 5, pp. 236–245, 2019.

[22] Liu et al., “Detecting cancer metastases on gigapixel pathology images,” arXiv preprint arXiv:1703.02442, 2017.

[23] James L Bennington, “Cancer of the kidney—etiology, epidemiology, and pathology,” Cancer, vol. 32, no. 5, pp. 1017– 1029, 1973.

[24] Grignon et al., “Clear cell renal cell carcinoma,” Clinics in Laboratory Medicine, vol. 25, no. 2, pp. 305–316, 2005.

[25] Akhtar et al., “Papillary renal cell carcinoma (prcc): an update,” Advances in Anatomic Pathology, vol. 26, no. 2, pp. 124–132, 2019.

[26] Vera-Badillo et al., “Chromophobe renal cell carcinoma: a review of an uncommon entity,” International Journal of Urology, vol. 19, no. 10, pp. 894–900, 2012.

[27] Wen et al., “A discriminative feature learning approach for deep face recognition,” in European Conference on Computer Vision. Springer, 2016, pp. 499–515.

[28] He et al., “Deep residual learning for image recognition,” in Proceedings of the IEEE Conference on Computer Vision and Pattern Recognition, 2016, pp. 770–778.

[29] Deng et al., “Imagenet: A large-scale hierarchical image database,” in 2009 IEEE Conference on Computer Vision and Pattern Recognition, 2009, pp. 248–255.

[30] Naylor et al., “Segmentation of nuclei in histopathology images by deep regression of the distance map,” IEEE Transactions on Medical Imaging, vol. 38, no. 2, pp. 448–459, 2018.

[31] Abdeltawab et al., “A deep learning framework for automated classification of histopathological kidney whole-slide images,” Journal of Pathology Informatics, vol. 13, pp. 100093, 2022.

[32] X. Li, C. Li, M.M. Rahaman, et al., “A comprehensive review of computer-aided whole-slide image analysis: from datasets to feature extraction, segmentation, classification and detection approaches,” Artificial Intelligence Review, vol. 55, pp. 4809– 4878, 2022.

[33] Pin et al., “Automatic cell nuclei segmentation and classification of breast cancer histopathology images,” Signal Processing, vol. 122, pp. 1–13, 2016.

[34] Q. Kang, Q. Lao, and T. Fevens, Nuclei Segmentation in Histopathological Images Using Two-Stage Learning, vol. 11764 of Lecture Notes in Computer Science, Springer, Cham, 2019.

[35] Martin Wattenberg, Fernanda Viégas, and Ian Johnson, “How to use t-sne effectively,” Distill, vol. 1, no. 10, pp. e2, 2016.

